# A new Mendelian Randomization method to estimate causal effects of multivariable brain imaging exposures

**DOI:** 10.1101/2021.10.01.462221

**Authors:** Chen Mo, Zhenyao Ye, Hongjie Ke, Tong Lu, Travis Canida, Song Liu, Qiong Wu, Zhiwei Zhao, Yizhou Ma, L. Elliot Hong, Peter Kochunov, Tianzhou Ma, Shuo Chen

## Abstract

The advent of simultaneously collected imaging-genetics data in large study cohorts provides an unprecedented opportunity to assess the causal effect of brain imaging traits on externally measured experimental results (e.g., cognitive tests) by treating genetic variants as instrumental variables. However, classic Mendelian Randomization methods are limited when handling high-throughput imaging traits as exposures to identify causal effects. We propose a new Mendelian Randomization framework to jointly select instrumental variables and imaging exposures, and then estimate the causal effect of multivariable imaging data on the outcome. We validate the proposed method with extensive data analyses and compare it with existing methods. We further apply our method to evaluate the causal effect of white matter microstructure integrity on cognitive function. The findings suggest that our method achieved better performance regarding sensitivity, bias, and false discovery rate compared to individually assessing the causal effect of a single exposure and jointly assessing the causal effect of multiple exposures without dimension reduction. Our application results indicated that WM measures across different tracts have a joint causal effect that significantly impacts the cognitive function among the participants from the UK Biobank.

## 1. Introduction

Imaging genetics is an emerging field that combines genetic and multi-modal brain imaging data to investigate the genetic effects on brain function or structure and to understand the neurogenetic mechanism of mental and neurological disorders and related disease and behavior phenotypes. Previous studies have used imaging genetics approaches to cognition, behavior in health and complex diseases.^1–6^ One increasingly important goal of imaging genetics studies is to test for causal imaging features on disease and related outcomes; and scalable methods to target this goal are in urgent need.^7,8^

Mendelian randomization (MR) methods estimate the causal effect of a modifiable exposure on an outcome in an observational study by employing genetic variants as instrumental variables (IVs).^9–11^ They address the limitations of traditional observational epidemiology regarding unobservable confounding and reverse causation^10,12–14^ and have been widely used in studies of potential causal inference.^15,16^ To successfully examine the causal effect, three key IV assumptions need to be met for MR analyses: i) IVs must be associated with the exposure of interest; ii) IVs must not be associated with confounders of the exposure-outcome association; and, iii) IVs must not affect the outcome except possibly through the exposure variable.^10,17–19^ MR experiments have generally relied on genetic variants associated with a single exposure to avoid violations of IV assumptions (ii) and (iii). However in practice, most variants are pleiotropic and associated with multiple exposures that cannot be ignored.^11^

Fig 1A shows the classical MR framework with multiple IVs and only one single imaging exposure. The classical MR method, such as the inverse-variance-weighted (IVW) approach, can estimate the causal effect of individual exposures using valid IVs following the fixed effect meta-analysis.^20^ However, especially in neuroimaging studies, MR analyses on only one imaging trait fail to completely capture the causal effects because these kind of analyses ignore the impact from other imaging traits, given that imaging traits are highly correlated. In addition, the presence of pleiotropic genetic variants will ultimately lead to inflated type I error rates and inadequate statistical power in MR analyses. For example, in Fig 1B, imaging traits have complex interconnections and may result in a combined effect coming from multiple traits rather than from a single exposure on the outcome. Their spatial dependency has created a few analytical challenges. Firstly, existing MR methods for multiple exposures allowed us to estimate causal effects of different exposures simultaneously on outcome, assuming additive effects.^7,11,21^ However, these methods are restricted by complicated horizontal pleiotropy and multicollinearity when the exposures are highly correlated as in the case of imaging features.^22^ Specifically, increasing the number of IVs and exposures makes the validation of IV assumptions challenging, consequently leading to biased causal estimates and false-positive causal relationships.^11^ Secondly, the framework involves multiple IVs and imaging exposures and usually cannot specify the subset of IVs with its influenced exposures, while preserving the validity of IV assumptions for all. Therefore, it becomes increasingly important to identify the subsets of strongly associated IVs and exposures as guided by their causal relationship with the outcome

**Fig. 1.**
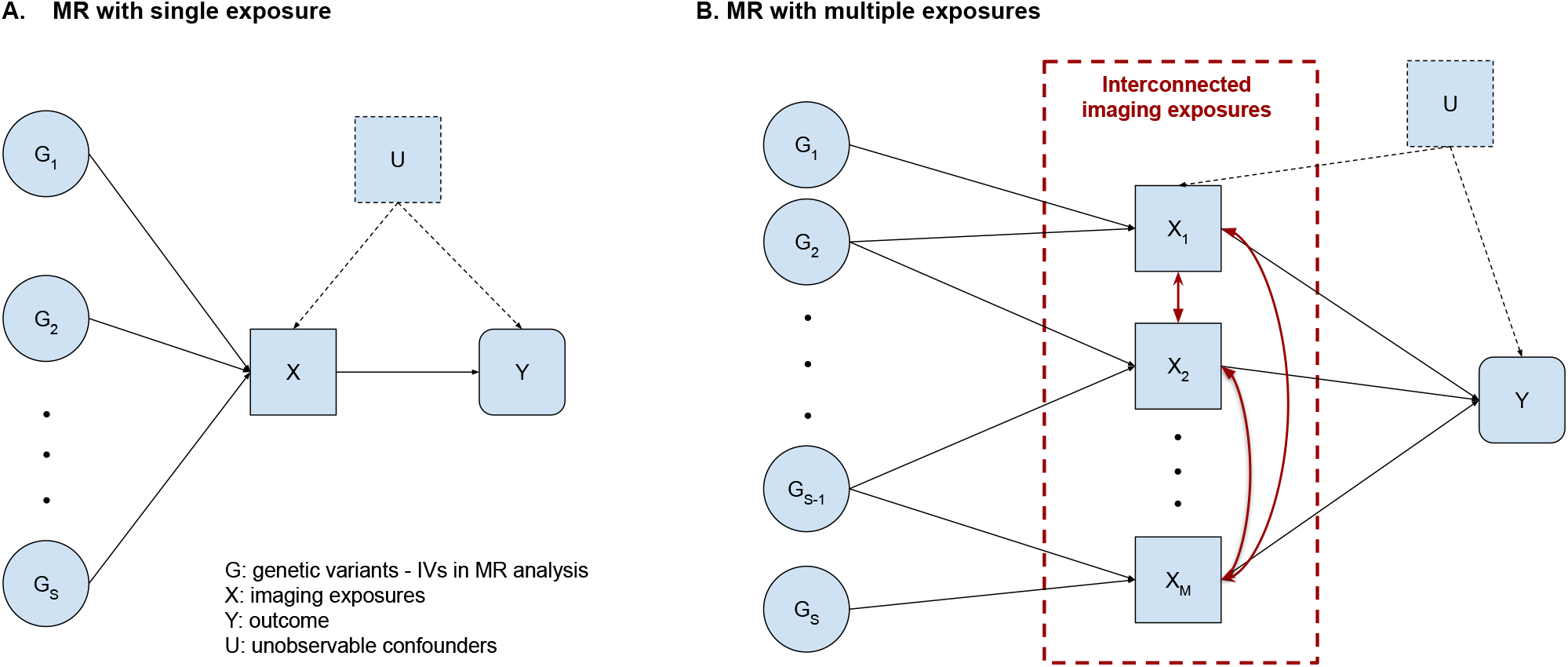
Mendelian Randomization with a single exposure (left) and multiple dependent exposures (right).

We propose a new method to address the aforementioned issues in MR analyses on multiple imaging exposures. Our method primarily selects a set of exposures that share a common set of IVs guided by data-driven submatrix identification algorithms.^23,24^ This method integrates the most informative features from exposures while reducing the burden of horizontal pleiotropy introduced by including too many exposures and IVs simultaneously in the MR model. In this study, we illustrated the application of our method using data from the UK Biobank (UKB) to examine the causal effects of white matter microstructure integrity (WM) measured with factional anisotropy (FA) on cognitive function. We also carried out simulation studies to compare the proposed method with existing MR methods. Both the application and simulation results demonstrated improved causality estimation. In this initial work, we focus on the individual-level data in one-sample MR analysis. We provide a detailed introduction of our method in section 2, the application to UKB data in section 3, simulation studies in section 4, and conclude with a comprehensive discussion in section 5.

## 2. Methods

In our application, brain imaging variables are multivariable exposures in the MR analysis (see Fig 1). The high-dimensionality of exposure variables leads to two new challenges: i) identifying causal exposures and corresponding IVs and ii) causal effect estimation for dependent multivariable exposures. Specifically, it is challenging to identify a subset of imaging variables with causal effects on the outcome and more importantly to extract a set of IVs that are jointly valid for the selected imaging exposures. To address these issues, we estimate the integrative causal effects of a set of dependent imaging exposures on the outcome. We provide an overview of our three-step approach and elaborate the procedures in the following subsections.

Our goal is to simultaneously select causal imaging exposures and corresponding valid IVs, such that each selected imaging exposure has causal effect based on the selected IVs. At the same time, each selected genetic variant is a valid IV for all selected imaging exposure IVs, satisfying the three commonly assumptions in MR analysis. Therefore, the IV set and exposure set selection procedures are interactive and can be subject to substantial false positive and false negative errors using an iterative procedure. We propose a new objective function for joint IV and exposure set selection.

### 2.1. Step 1 : Mendelian randomization analysis on a single imaging exposure

We first perform MR analysis on each imaging exposure with loci of interest and assess the validity of IVs by following the guideline for MR investigations proposed by Burgess et al. (2019).^19^ We record the validity as a indicator function for each pair of genetic variant and imaging exposure *a*_*sm*_ in matrix ***A***_***S×M***_ = {*a*_*sm*_}. Similarly, we store the single-exposure MR analysis results in a matrix ***Q***_***S×M***_ = {*q*_*sm*_}. We have

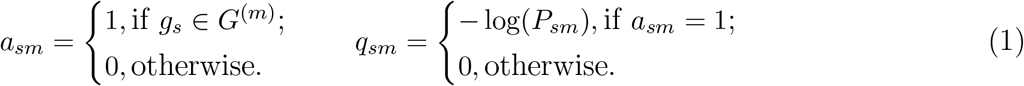

### 2.2. Step 2: Joint instrumental variables and imaging exposures selection

Next, we detect a submatrix from a large matrix of genetic variants and imaging exposures ***W*** = ***A*** ∘ ***Q***, where ∘ is the Hadamard product. Our objective function is an, *l*_0_ shrinkage function to extract the maximal number of valid imaging exposure-IV pairs with minimally sized IV set and imaging variable set. Specifically:

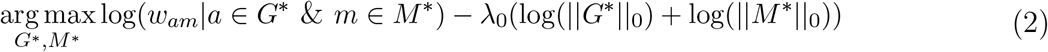

where *G*^*^ is the IV set and *M* ^*^ is the imaging exposure set, ||||_0_ is the cardinality measure of a set, and *λ*_0_ is a tuning parameter. The first item ensures the maximal information can be included based on selected *G*^*^ and *M* ^*^, while the second term penalizes the cardinality of *G*^*^ and *M* ^*^ to avoid the false positive errors. The objective function is non-convex due to the, *l*_0_ term, and thus computationally intensive. We employ greedy algorithms to implement the objective function for large-sized ***W*** ^23^ and exhausting search algorithms for medium-small ***W*** .^24^ Both algorithms can be conveniently extended to multiple sets of *G*^*^ and *M* ^*^.

Specifically, we can search the optimal submatrix ***W*** ^*^ determined by *M* ^*^ and *G*^*^ that contains the most informative features following a general iterative procedure:

1. Find a submatrix 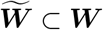 by a greedy search algorithm^23^ to approximately maximize the objective function.
2. Subtract the average of 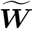 from each of its entries in ***W***.
3. repeat until convergence criteria is met.

This algorithm searches the solution of the objective function in an iterative-residual fashion, which captures the most informative features of the data matrix (***W***) that are of potential causal effect inference^24^ with parsimonious IV set and exposure set *M* ^*^ and *G*^*^.

### 2.3. Step 3: Causal effect identification for multiple imaging exposures

Given a common IV set *G*^*^ and a set of imaging exposures *M* ^*^, we attempt to estimate the causal effect of multiple dependent exposures through MR analysis. It is challenging to identify the causal effects of imaging exposures because the highly correlated exposures can lead to imprecise causal effect estimation.^25^ This is a common issue that mediation analysis has been facing.^26^ We adopt commonly used statistical techniques in imaging causal mediation analysis to transform the imaging exposures into a set of orthogonal variables. Let **M**^*^ = (**M**_1_, … **M**_*n*_)^T^ ∈ ℝ^*n*×*p*^ denote the matrix for *p* selected imaging features across *n* subjects, and 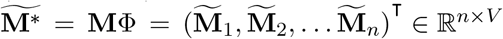 be the matrix of orthogonally transformed imaging variables where Φ ∈ ℝ^*p*×*V*^ is the transforming matrix. We can estimate Φ and 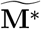 based on the procedure described in Chén et al. (2018).^26^ Furthermore, we can implement sparsity on loadings for the components to improve the interpretability.^27^

We next perform MR analysis on *V* orthogonal imaging factors Φ with independent causal effects. Given these conditions, we only need the MR analysis on individual factors because the orthogonal imaging factors only have additive causal effects. For an imaging factor 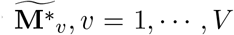, we can estimate its causal effect on the outcome (*Y*) using uncorrelated IVs (*G*^*^ = {*g*_1_, …, *g*_*S*_} ⊆ *G*) through the IVW method as follows:

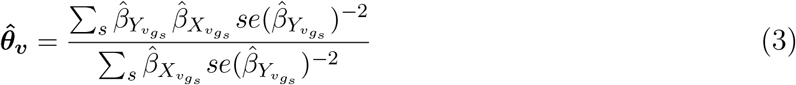

where 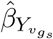 and 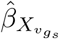 are the genetic associations based on the regression of the outcome (*Y*) and the imaging factor 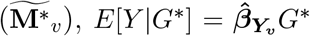 and 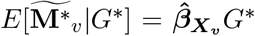, respectively, with the approximated standard error 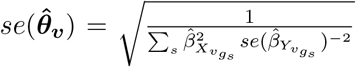 summing across the estimates from all IVs in *G*^*^.^20^ The overall causal effect of all exposures given the identified imaging factors can be simply expressed as 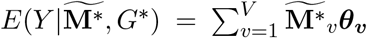 In case that the IVs are correlated, the IVW can be extended to account for the correlation matrix using methods such as the generalized weighted linear regression,^22,28^ Causal Direction-Ratio,^29^ and Causal Direction-Egger.^29^ We leave the details of using correlated IVs in the future study and focus on uncorrelated IVs in our current study.

#### Remarks

Our MR framework consists of three steps as follows: step 1: select IV candidates associated with each imaging exposures; step 2: extract submatrices of valid IVs and corresponding imaging exposures; step 3: conduct MR analysis based on IVs and transformed imaging exposures in the extracted submatrices.

## 3. Application to evaluate the causal effect of white matter microstructure integrity on cognitive function

### 3.1. Data and study cohort

We applied our new method to a sample of 35,291 unrelated participants (white ethnicity backgrounds aged 40-69) extracted from the UKB to evaluate causal effect of white matter integrity on cognitive functions.^30^ The exposures consisted of forty regional brain FA measures derived from diffusion MRI based on the preprocessing workflow of the Enhancing Neuro Imaging Genetics Meta Analysis (ENIGMA) consortium.^31^ The outcome was the intelligence ***g*** estimated from five cognitive traits related to the following four domains: processing speed, perceptual reasoning, executive function and fluid intelligence.

The intelligence ***g*** was estimated among 10,979 participants with cognitive data. The missing values were substituted by the average of imputed values based on predictive mean matching (PMM) method implemented in *R* package *mice (v3.13.0)*.^32^ We estimated this latent general intelligence factor accounting for 59% of the total variance of the cognitive traits using R package *psych (v 2.1.9)*.^33^

The genotypic data was available for all participants involved in the analysis. We implemented quality control with following inclusive thresholds: minor allele frequency (MAF) > 0.01, Hardy-Weinberg equilibrium (HWE) > 0.001, missingness per marker (GENO) < 0.05, and missingness per individual (MIND) < 0.02 by PLINK (v1.9).^34^ We removed highly correlated genetic variants (*r*^2^ < 0.5) via LD clumping and used the variants in gene VCAN as potential IVs since many studies have discovered significant associations between VCAN and the FA measures, as listed in the NHGRI-EBI GWAS Catalog.^35^ We adjusted for variables such as sex, age, body mass index (BMI), genotyping chip type and top ten PCs of population admixture in our MR analysis.

### 3.2. Results

We identified 31 out of 40 FA measures having a significant association (*p*-value < 0.05 adjusted with false discovery rate^36^) with intelligence ***g*** after data preprocessing. In total, we found 27 genetic variants in *VCAN* that had highly significant associations (*p*-value < 5 × 10^−7^) with at least one of the 31 FA measures. These variants were weakly correlated with each other. As shown in Fig 2B, the heatmap presented the causal effect significance (− log(*p*-value)) estimated from MR using a single IV for every exposure, given the rows and columns represented the 27 IVs and 31 FAs, respectively.

**Fig. 2.**
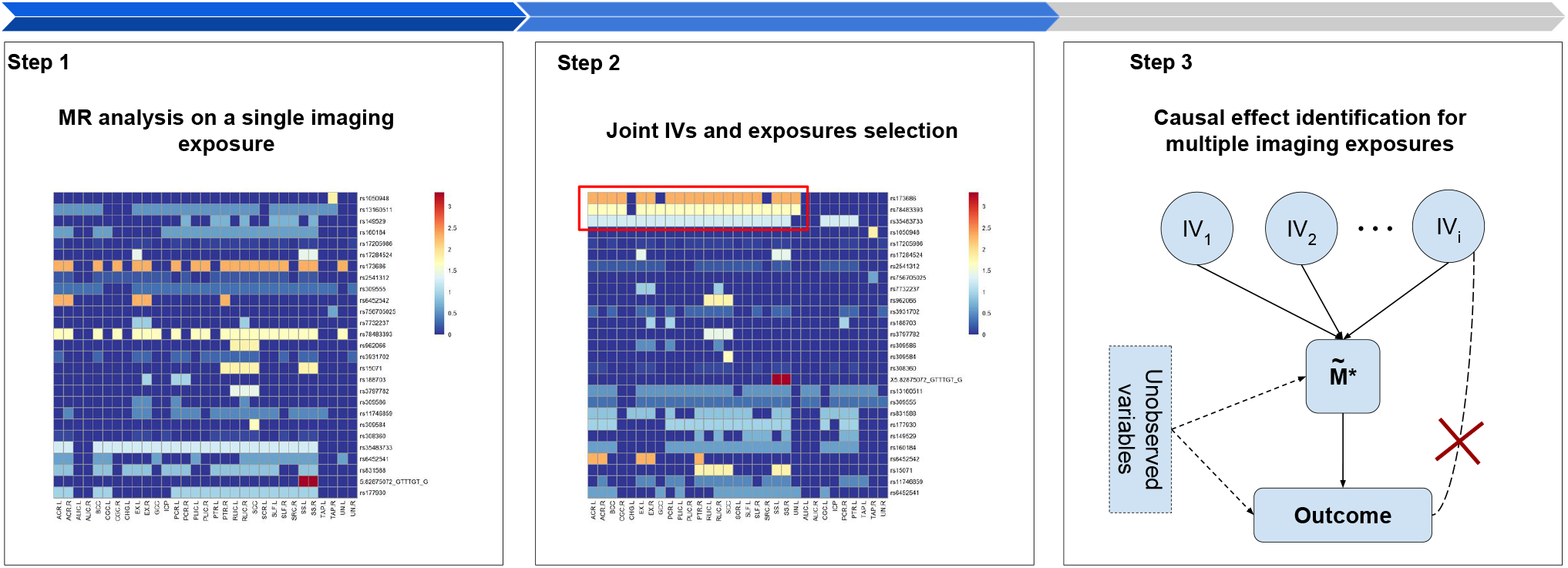
Overview of analysis framework. Our MR analysis method consists of three main steps. The heatmap (left) shows the raw unorganized matrix of −*logP*-value in the first analysis step; the heatmap (middle) shows the matrix after submatrix identification in the second step, showing a cluster of most informative features; and, the diagram (right) shows the MR analysis on the identified features with selected IVs in the last step.

We observed that FAs affected intelligence ***g*** in different levels and some of these measures had similar effects based on their common IVs (see heatmap (left) Fig 2), although they were arranged in a random order. We further detected an informative cluster consisting of 22 FA measures with 3 common IVs in this unorganized structure by implementing our objective function. These FA measures were: bilateral anterior corona radiata, body of corpus callosum, cingulum cingulate gyrus (right), cingulum hippocampus (left), bilateral external capsule, genu of corpus callosum, posterior corona radiata (left), bilateral posterior limb of internal capsule, posterior thalamic radiation (right), bilateral retrolenticular part of internal capsule, splenium of corpus callosum, bilateral superior corona radiata, bilateral superior longitudinal fasciculus, bilateral sagittal stratum, and uncinate fasciculus (left). The 3 SNPs of VCAN on chromosome 5 included: rs173686, rs35483733, rs78483393, having reported association with white matter integrity in the previous study.^37^ Fig 3B (upper) showed the correlation matrix of the 22 FA measures selected. These imaging exposures had moderate to high correlations between each other, suggesting non-identifiable causal effects based on existing MR methods with independent causal effects. Following step 3 in our method, we transformed the 22 FA measures into a single general factor. The general factor of FA (*g*FA) was estimated based on 3 components achieving the highest percentage (59%) of common variance among 22 FA measures. The number of components used was approximated by a parallel analysis (see Fig 3B (lower)). The loading for the rest factors was unstable based on bootstrap model validation, and thus were not used.

**Fig. 3.**
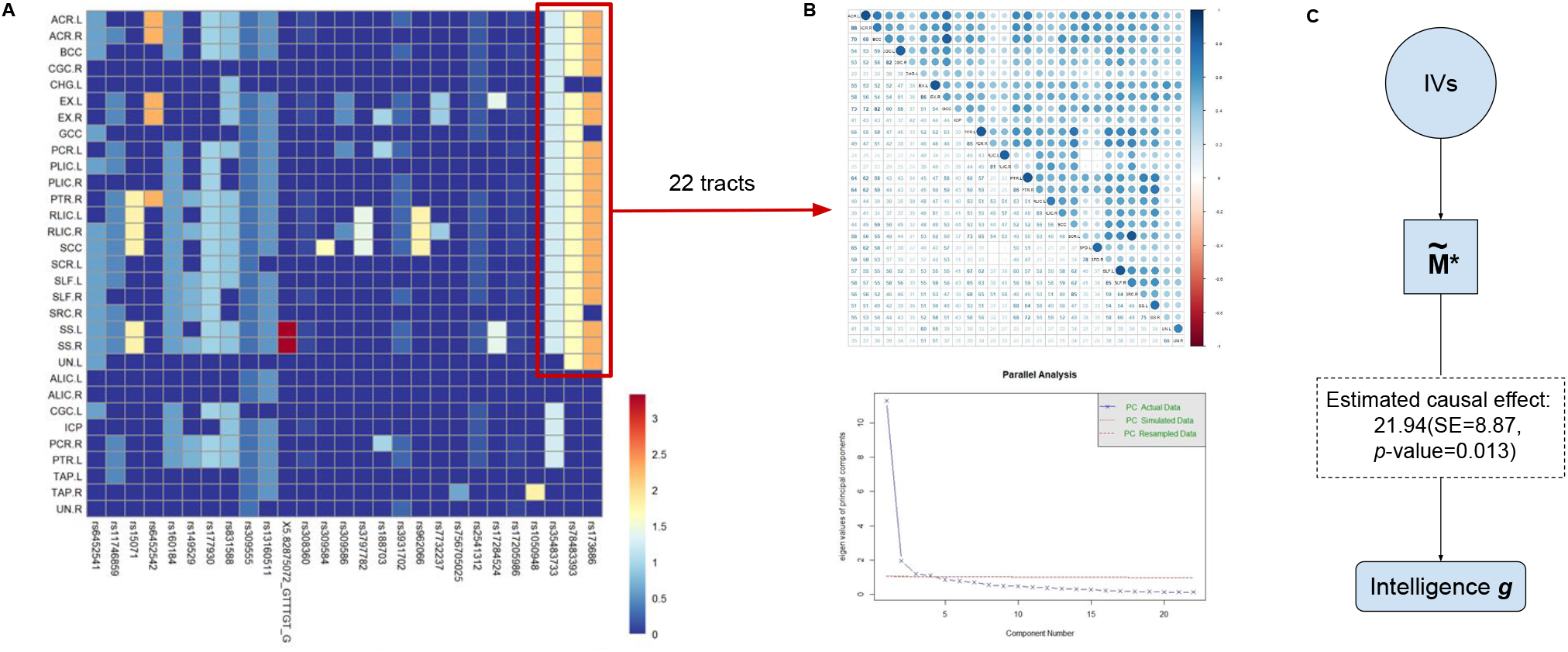
Mendelian randomization analysis results of imaging exposures and cognitive function. A shows the 22 FA tracts identified within a submatrix extracted from 31 FA tracts. The lowest significance was shown in dark blue whereas red indicated the highest significance of causal effect; B shows the matrix of pair-wise correlation matrix of the 22 tracts along with their parallel analysis based on PCA for estimating orthogonal factors; and, C shows the MR analysis with its final results of the causal effect across the uncorrelated orthogonal imaging factors.

Next, we assessed the comprehensive causal effect of FAs (*g*FA) on cognitive function (intelligence ***g***) via classical IVW-based MR method using the *MendelianRandomization (v0.5.1)* package in R.^38^ The results revealed that *g*FA had significant causal effect on intelligence ***g*** (*β* = 21.94, *SE*_*β*_ = 8.87, *p*-value = 0.013). We also explored the causal effect estimated via MR methods incorporating penalized regression,^39^ robust regression,^40^ and leave-one-out^41^ to assess the consistency of the causal estimates and possible IV outliers. The results were all consistent with the classical IVW method showing a significant causal effect of *g*FA on intelligence ***g***. All in agreement, these results consistently revealed that the increase of white matter microstructure integrity can cause the improvement of performance regarding cognitive function tests.

## 4. Simulation

We carried out simulation studies to evaluate our proposed framework of MR analysis for quantitative traits under the one-sample case. For *n* = 500 individuals, we first randomly simulated genotypes **X**_500×20_ for 20 uncorrelated genetic variants (i.e. IVs in the MR analyses). Here we assumed there was an underlying true factor of imaging exposure *Mf*_500×1_ = **X***α*_20×1_, where *α*^*T*^ = (2, 2, …, 2, 0, 0, …, 0) measured the effect that genetic variants had on exposures. We also assumed that only 10 simulated genetic variants had true effect on this underlying exposure factor, whereas the other 10 variants had no true effect. Next, we generated 20 observed imaging exposures with true casual effects on the outcome by 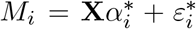, where 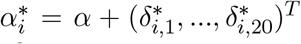 and 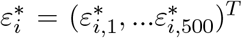. In addition, we simulated another 20 observed imaging exposures without true casual effects on the outcome by 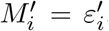, where 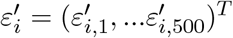. Here 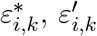 and 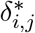 are all *i.i.d* random noise with standard normal distribution, where *i, j* ∈ {1, …, 20} and *k* ∈ {1, …, 500}. Finally, we simulated the outcome data using the true exposure factor, i.e. *Y*_500×1_ = *β* * *Mf* + *ε*_500×1_, and *ε* = (*ε*_1_, …, *ε*_500_)^*T*^ is another set of standard normal random noises. We consider two cases for the causal effect size: large (*β* = 1) and small (*β* = 0.5).

Under this simulation setting, three types of MR analyses were implemented and their performances were compared. The first one was our method, which implemented LAS^24^ to identify submatrices before MR and only included a subset of essential imaging exposures in the MR model. The second method included all 40 imaging exposures in the MR model, and the third one simply ran 40 MR models with single exposure independently. To evaluate, we calculated the bias of the point estimates for causal effect *β*, and the sensitivity and False Discovery Rate (FDR) for correctly selecting the true imaging exposures with casual effect. For the later part, the second method just simply included all the exposures (i.e. selecting all) and the third method made the selection based on its p-values with a Benjamini-Hochberg correction (number of comparisons is 800).

We ran the simulation for 500 replications, and the results are given in Table 1. The computation time was 60 seconds per replication using a desktop with CPU 3.40GHz and RAM 64GB on average. In both the large effect and small effect settings, our method achieved smaller bias in estimating the causal effect compared to the method using all 40 imaging exposures (0.108 vs. 0.924 for large effect, 0.05 vs. 0.473 for small effect). In terms of the selection of causal imaging exposures, our method had substantially decreased FDR (0.15 and 0.148) while still maintaining a sensitivity closed to 1 (0.947 and 0.945).

**Table 1.** Simulation results for two different causal effects size *β* = 1 and *β* = 0.

| Method | Simulation results with $\beta = 1$ | | |
| --- | --- | --- | --- |
| | Bias of $\hat{\beta}$ | Sensitivity | FDR |
| MR with exposures selected (our method) | 0.108 (0.084) | 0.947 (0.075) | 0.15 (0.157) |
| MR with all exposures | 0.924 (0.213) | 1 (0) | 0.5 (0) |
| MR with a single exposure | - | 1 (0) | 0.5 (0) |
| Method | Simulation results with $\beta = 0.5$ | | |
| | Bias of $\hat{\beta}$ | Sensitivity | FDR |
| MR with exposures selected (our method) | 0.05 (0.045) | 0.945 (0.077) | 0.148 (0.155) |
| MR with all exposures | 0.473 (0.107) | 1 (0) | 0.5 (0) |
| MR with a single exposure | - | 1 (0) | 0.5 (0) |

## 5. Discussion

We developed a new MR framework to evaluate the causal effects of inter-correlated mutlivariable brain imaging exposures on outcomes. Our approach provides a viable solution to estimate the causal effect of objectively measured characteristics of the central nervous system on externally measured neuropsychological test results by leveraging imaging-genetics data. The utility of genetic variants as instrumental variables leads to unbiased estimates of causal effects free from confounding effects from numerous environmental factors.

The MR analysis with brain imaging variables as exposures is intrinsically challenging. The selection of valid IVs for all imaging exposures and the selection of causal imaging exposures are complex and numerically difficult. We propose a new objective function to select exposures and IVs for maximal information while controlling false positive error rate by penalizing the cardinality of IV and imaging sets. The selected imaging variables provide spatially-specific causes for the externally measured test results. The shared IV set also becomes the foundation to transform the imaging exposures to orthogonal and causal independent factors as the IVs are valid for any of the imaging variables. Last, we estimate the causal effects of the transformed exposures of selected imaging variables and make inference.

Compared to previous studies that only repeatedly tested the associations between white matter microstructure integrity and cognitive function, our analysis revealed a significant comprehensive *causal* relationship between them. The decrease of white matter microstructure integrity causes the decline in cognitive function while adjusting for age, sex and other covariates mentioned above. Although our current analyses focus on region-level imaging variables, our method can be extended to voxel-level analyses. We also assume that there exists no cyclic causal effects between multiple exposures and the outcome. Our study aims to address issues of the multiple-exposure MR particularly in the one-sample studies because the existing studies and resources of summary statistics for all exposures included are restricted and more difficult to ensure valid IVs to achieve two-sample scenario.

In summary, our MR analysis framework with multivariable imaging exposures opens a new avenue for imaging-genetics data analysis and causal inference. This study currently focuses on MR analysis using uncorrelated IVs. Our framework can also be extended to MR analysis using correlated IVs adopting the new MR methods that account for complex covariance structure among IVs in future studies.

## Acknowledgements

Preprint of an article published in Pacific Symposium on Biocomputing © 2021 World Scientific Publishing Co., Singapore, http://psb.stanford.edu/.

## Availability of data and materials

The data used in this study are available in the UK Biobank, https://www.ukbiobank.ac.uk/. We provide the GWAS summary statistics and codes in the GitHub repository, https://github.com/kehongjie/ImagingMR.

## Authors’ contributions

CM, HK, ZY, SC and TM developed the method and wrote the manuscript. CM, HK and ZY performed the analysis and results visualization. SC and TM supervised the project and provided conceptualization. TL, TC, SL, QW, ZZ, YM, LEH and PK contributed to manuscript editing and provided critical feedback and help to shape the research, analysis and manuscript. All authors have read and approved the manuscript.

